# Electrophysiology reveals the neural dynamics of naturalistic auditory language processing: event-related potentials reflect continuous model updates

**DOI:** 10.1101/062299

**Authors:** Phillip M. Alday, Matthias Schlesewsky, Ina Bornkessel-Schlesewsky

## Abstract

The recent trend away from ANOVA-based analyses places experimental investigations into the neurobiology of cognition in more naturalistic and ecologically valid designs within reach. Using mixed-effects models for epoch-based regression, we demonstrate the feasibility of examining event-related potentials (ERPs), and in particular the N400, to study the neural dynamics of auditory language processing in a naturalistic setting. Despite the large variability between trials during naturalistic stimulation, we replicated previous findings from the literature: frequency, animacy, word order. This suggests a new perspective on ERPs, namely as a continuous modulation reflecting continuous model updates (cf. Friston, 2005) instead of a series of discrete and essentially sequential processes.

## 1. Introduction

In real-life situations, the human brain is routinely confronted with complex, continuous and multimodal sensory input. Such natural stimulation differs strikingly from traditional laboratory settings, in which test subjects are presented with controlled, impoverished and often isolated stimuli (e.g. individual pictures or words) and often perform artificial tasks. Accordingly, cognitive neuroscience has seen an increasing trend towards more naturalistic experimental paradigms (Hasson and Honey, 2012), in which complex, dynamic stimuli (e.g. movies, natural stories) are presented without an explicit task (e.g. Hasson et al., 2004, Hasson 2008; Skipper et al., 2009; Whitney et al., 2009; Brennan et al., 2012; Lerner et al., 2011; Conroy et al., 2013; Hanke et al., 2014).

In spite of being uncontrolled, naturalistic stimuli have been shown to engender distinctive and reliable patterns of brain activity (Hasson et al., 2010). However, they also pose unique challenges with respect to data analysis (e.g. Hasson and Honey, 2012, cf. also the 2014 Real-life neural processing contest, in which researchers were invited to develop novel analysis techniques for brain imaging data obtained using complex, naturalistic stimulation). To date, the discussion of these challenges has focused primarily on neuroimaging data and, in the majority of cases, on visual stimulation. Naturalistic stimuli in the auditory modality, by contrast, give rise to an additional and unique set of problems, particularly when examined using techniques with a high temporal resolution such as electroencephalography (EEG) or magnetoencephalography (MEG). Consider the case of language processing: in contrast to typical, controlled laboratory stimuli, a natural story or dialogue contains words that vary vastly in length, a stimulus property to which EEG and MEG are particularly sensitive because of their superb temporal resolution. The characteristic unfolding over time of auditory stimuli is already evident when evoked electrophysiological responses are compared in more traditional, controlled studies — the endogenous components show increased latency and a broader temporal distribution (see for example Wolff et al., 2008, where the same study was carried out in the auditory and visual modalities). EEG and MEG studies with naturalistic stimuli consequently tend to use the so less naturalistic visual modality (segmented, rapid-serial visual presentation, e.g. Frank et al. (2015); or natural reading combined with eye-tracking, e.g. Kretzschmar et al. (2013); Hutzler et al. (2007)).

Given current data-analysis techniques, these distinctive properties of the auditory modality impose severe limitations on our ability to conduct and interpret naturalistic auditory experiments, particularly when seeking to address questions related to time course information in the range of tens - or even hundreds - of milliseconds. Here, we present a new synthesis of analysis techniques that addresses this problem using linear mixed-effects modeling. We further provide an initial demonstration of the feasibility of this approach for studying auditorily presented naturalistic stimuli using electrophysiology.

For this initial exploratory study, we focus on the N400 event-related potential (ERP), a negative potential deflection with a centro-parietal maximum and a peak latency of approximately 400 ms, but the methodology should apply to other ERP components as well.

### 1.1 *The N400*

The N400 is well suited to the purposes of the present study, since it is highly robust and possibly the most researched ERP component in the neurobiology of language (see Kutas and Federmeier, 2011, for a recent review). Although the exact neurocognitive mechanism(s) that the N400 indexes are still under debate, it can be broadly described as being sensitive to manipulations of expectation and its fulfillment (cf. Kutas and Federmeier, 2000, Kutas 2011; Lotze et al., 2011; Lau et al., 2008; Hagoort, 2007). This can be seen most clearly in the sensitivity of the N400 to word frequency, cloze probability and contextual constraint, but also to manipulations of more complex linguistic cues such as animacy, word order and morphological case as well as the interaction of these factors (Bornkessel and Schlesewsky, 2006; Bornkessel-Schlesewsky and Schlesewsky, 2009). Importantly for the examination of naturalistic stimuli, N400 amplitude is known to vary parametrically with modulations of these cues, thus making it well suited to modeling neural activity based on continuous predictors and activity fluctuations on a trial-by-trial basis (cf. Cummings et al., 2006; Roehm et al., 2013; Sassenhagen et al., 2014; Payne et al., 2015).

More recently, researchers have attempted to quantify expectation using measures derived from information theory, such as surprisal. These have enjoyed some success as a parsing oracle in computational psycholinguistics (Hale, 2001; Levy, 2008; cf. Smith and Levy, 2013, for a computational approach applied to eye-tracking data) and have been shown to correlate with N400 amplitude for naturalistic stimuli (real sentences taken from an eye-tracking corpus) presented with RSVP (Frank et al., 2015).

### 1.2 *Mixed-effects Models*

Mixed-effects models present several advantages over traditional repeated-measures ANOVA for the exploration presented here. First, they yield quantitative results, estimating the actual difference between conditions instead of merely the significance of the difference. Second, they can easily accommodate both quantitative and qualitative independent variables, allowing us to integrate measures such as frequency without relying on dichotomization and the associated loss of power (cf. MacCallum et al., 2002) Finally, they are better able to accommodate unbalanced designs than traditional ANOVA methods. A more comprehensive review of mixed-effects models, especially as it pertains to the study at hand, can be found in the supplementary materials. For a similar approach at the sentence-1 level, we refer the interested reader to Payne et al. (2015), which also includes a review of mixed-effect modelling in its supplementary materials, albeit with a somewhat different focus.

## 2. Materials and methods

### 2.1 *Participants*

Fifty-seven right-handed, monolingually raised, German native speakers with normal hearing, mostly students at the Universities of Marburg and of Mainz participated in the present study after giving written informed consent. The experiment was performed in accordance with the ethical standards laid down in the Declaration of Helsinki. Approval by an ethics review board was not required, as current regulations in Germany specify that ethics approval for ERP experiments is only required when participants are patients, children or older adults (> 65 years) (?). Three subjects were eliminated due to technical issues, one for psychotropic medication, and one for excessive yawning, leaving a total of 52 subjects (mean age 24.2 std.dev 2.55; 32 women) for the final analysis.

### 2.2 *Experimental stimulus and procedure*

Participants listened passively to a story roughly 23 minutes in length while looking at a fixation star. Subjects were instructed to blink as little as possible, but that it was better to blink than to tense up from discomfort. After the auditory presentation, test subjects filled out a short comprehension questionnaire to control for attentiveness.

The story recording, a slightly modified version of the German novella “Der Kuli Klimgun” by Max Dauthendey read by a trained male native speaker of German, was previously used in an fMRI study by Whitney et al. (2009). For each word in the transcribed text, a lingusitically trained native speaker of German provided an annotation for the prominence features “animacy”, “morphological case marking” (morphological ambiguity was not resolved even if syntactically unambiguous), “definiteness” (i.e. whether the definite article “the” was present), “humanness” and “position” (initial or not for nominal arguments). Tags were placed at the position that the prominence information was “new”; an automated process created a duplicate tagging where the new information was repeated for the rest of its constituent phrase (e.g. copying case-marked from the determiner to the head noun). Absolute (“corpus”) frequency estimates were extracted programmatically from the Leipziger Wortschatz using the Python 3 update to libleipzig-python. Relative frequencies were calculated as the ratio of orthographic tokens to orthographic types.

### 2.3 *EEG recording and preprocessing*

EEG data were recorded from 27 Ag/AgCl electrodes fixed in an elastic cap (Easycap GmbH, Herrsching, Germany) using a BrainAmp amplifier (Brain Products GmbH, Gilching, Germany). Recordings were sampled at 500 Hz, referenced to the left mastoid and re-referenced to linked mastoids offline. All signal processing was performed using EEGLAB (Delorme and Makeig, 2004) and its accessary programs and plugins. Using sine-wave fitting, the EEG data were first cleaned of line noise (Clean-line plugin), and then automatically cleaned of artifacts using an programmatic procedure based upon ICA (MARA, Winkler et al., 2011). Although automatic procedures have come under some criticism for being both overly und insufficiently conservative in their selection (cf. Chaumon et al., 2015), they have the distinct advantage of being (nearly) deterministic and thus completely replicable as well as faster for large numbers of subjects, as in the present study. The majority of removed components were eye movements (blinks and saccades) as well as several with a single-electrode focus, generally lateralized. As the following analysis (see below) used electrodes exclusively on the centro-parietal midline, i.e. not lateral, the removal of these components is not problematic. The ICA decomposition was performed via Adaptive-Mixture ICA on data high-pass filtered at 1 Hz (to increase stationarity) and downsampled to 100Hz (for computational tractability) (Palmer et al., 2007) and backprojected onto the original data; no rank reduction was performed and as such 27 components were extracted. Subsequently, the original data were high-pass filtered at 0.3 Hz and 1682 segments extracted per test subject, time locked to the onset of content words (cf. “open-class words” in Payne et al., 2015; Van Petten and Kutas, 1991). This filter was chosen to remove slow signal drifts as traditional baselining makes little sense in the heterogeneous environment of naturalistic stimuli (cf. Frank et al., 2015, who additionally found that a heavier filter helped to remove correlation between the pre-stimulus and component time windows). All filtering was performed using EEGLAB’s `pop_eegfiltnew()` function.

### 2.4 *Data analysis*

We examined single trial mean amplitude in the time window 300-500ms, a typical time window for the N400 effect (Kutas and Federmeier, 2011; cf. Frank et al., 2015, Payne et al., 2015; see also Pernet et al., 2011; Bishop and Hardiman, 2010, for other single-trial analyses in traditional paradigms). To simplify the analysis, both computationally and in terms of comprehensibility, only data from the electrodes Cz, CPz, and Pz were used, following the centro-parietal distribution of the N400 (cf. Payne et al., 2015; see also the single-electrode analysis in Tremblay and Newman, 2015, for exploratory and demonstration purposes with generalized additive mixed-effects models). Single-trial epoch averages from these electrodes were analyzed using linear mixed-effects models (LMM, Pinheiro and Bates, 2000; Bates et al., 2015b, see supplementary materials for R session information, including package version of `lme4`).

### 2.5 *Statistical Methods*

For the analysis presented here, we use a minimal LMM with a single random-effects term for the intercept of the individual subjects. This is equivalent to assuming that all subjects react the same way to each experimental manipulation but may have different “baseline” activity. This is a plausible assumption for an initial exploration, where we focus less on interindividual variation and instead focus on the feasibility of measuring population-level effects across subjects. Furthermore, this is not in violation of Barr et al. (2013)’s advice, which is explicitly directed at *confirmative* studies. The reduced random-effects structure reduces the number of parameters to estimate, which (1) greatly increases the computational tractability of the exploration at hand and (2) allows us to focus the relatively low power of this experimental setup on the parameters of interest (cf. Bates et al., 2015a).

We omit a random-effect term for “item” as there are no “items” in the traditional psycholinguistic sense here (Clark, 1973). A random effect for “lexeme” is also not appropriate because while some lexemes appear multiple times (e.g., “Ali”, the name of the title character), many lexemes appear only once and this would lead to over parameterization.

No parameter for electrode was introduced into the model as this would have reduced overall power and increased computational complexity. The three electrodes used are close enough together that they should all have correlated and highly similar values, which means more data and thus more precise estimates. Mixed-effects models do not require that these measurements are explicitly averaged beforehand (complete pooling), but can use all measurements to provide better parameter estimates - in tuitively, the model “implicitly” averages the three measurements. The differences between electrodes become part of the residual error, but the extra information provided by additional measurements can nonetheless improve overall model fit. This also accommodates variation, due minor differences in physiology and cap placement between subjects better than a single-electrode analysis (cf. “optimized averaging” in Rousselet and Pernet, 2011).

Categorical variables were encoded with **sum encoding** (i.e. ANOVA-style coding), such that the model coefficient represents the size of the contrast from a given predictor level to the (grand) mean. For a two-level predictor, this is exactly half the difference between the two levels (because the mean is equidistant from both points).

As indicated above, the dependent measures is the single-trial average amplitude in the epoch from 300 to 500ms post stimulus onset.

For simpler models, we present the full model summary, including an estimation of the inter-subject variance and all estimated coefficients for the fixed effects, but for more complicated models, we present a contour plot of the effects as modelled (i.e. the predictions from the LMM) in the main text, with the full summary moved to the supplementary materials along with a brief selection of the strongest effects, as revealed by Type-II Wald F-tests (i.e. with `car::Anova()`, Fox and Weisberg, 2011). Type-II Wald tests have a number of problems (cf. Fox, 2016, pages 724-725, 737-738, and discussions on R-SIG-mixed-models), but even assuming that their results yield an anticonservative estimate, we can use them to get a rough impression of the overall effect structure (cf. Bolker et al., 2009). Model comparisons, or, more precisely, comparisons of model fit, were performed using the Akaike Information Criterion (AIC, Akaike, 1974), the Bayesian Information Criterion (BIC, Schwarz, 1978) and log-likelihood. AIC and BIC include a penalty for additional parameters and thus provide an integrated measure of fit and parsimony. For nested models, this comparison was performed as a likelihood-ratio test, but non-nested models lack a significance test for comparing fit. We do not include pseudo *R*^2^ values because these are problematic at best and misleading at worst in an LMM context (see supplementary materials).

For the model summaries, we view |*t*| > 2 (i.e., the estimate of the coefficient is more than twice as large as the error in the estimate) as being indicative of a reliable estimate in the sense that the estimate is distinguishable from noise. We view |*t*| < 2 as being unreliable estimates, which may be an indicator of low power or of a generally trivial effect. (We note that Baayen et al. (2008) use |*t*| > 2 as approximating the 5%-significance level.) For the Type-II Wald tests, we use the *p*-values as a rough indication of reliability of the estimate across all levels of a factor. This will become clearer with an example, and so we begin with a well-known modulator of the N400: frequency of a word in the language as a whole.

## 3. Results and Discussion

### 3.1 *Proof of Concept: Frequency*

In a natural story context, traditional ERP methodology with averaging and grand averaging yields waveforms that appear uninterpretable or even full of artifacts. From the perspective of continuous processing, this is not surprising at all. Some information is present before word onset via context (e.g. modifiers before a noun), which leads to ERPs that seem to show an effect very close to or even before zero. Some words are longer than others, which leads to a smearing of the traditional component structure, both at a single-trial and at the level of averages. These problems are clearly visible in Figure 1, which shows an ERP image (Jung et al., 2001) for a single participant for initial accusatives (roughly, an object-first word order), which are known to be dispreferred to initial nominatives (roughly, a subject-initial word order) (Schlesewsky et al., 2003) and thus should engender an N400 effect. However, a modulation of the ERP signal is nonetheless detectable in the N400 time window, indexing the processing of the new information available at the trigger point. As a proof of concept for our method, we first examine the well-established effect of frequency on N400 amplitude (see Kutas and Federmeier, 2011, for a review), the results of which are presented in Table 1.

#### 3.1.1 *Corpus Frequency*

The frequency of a word in the language as whole, *cor-pus frequency*, is known to correlate with N400 amplitude and to interact with cloze probability (see Kutas and Federmeier, 2011, for a review). Using the logarithmic frequency classes from the Leipzig Wortschatz, we can see in Table 1 that corpus frequency has a small, but reliable effect (only −0.06 *μ*V per frequency class, but *t* < −13 in the N400 time window). This is exactly what the literature predicts - frequency is not dominant in context-rich environments, but plays a distinct role (cf. Kutas and Federmeier, 2011; Dambacher et al., 2006).

Moreover, corpus frequency is insensitive to context as it represents global and not local information. Adding index, i.e. the ordinal position in the story, to the corpus frequency model does not improve it (minimal change in log-likelihood, no change in AIC, worsening of BIC, see Table S2 in the supplementary materials for the full comparison). This lack of improvement reflects the context insensitivity of corpus frequency, which is a global measure not dependent on the story context. (At the sentence level, there is evidence that ordinal position modulates the role of frequency, e.g. Van Petten and Kutas (1990), Payne et al. (2015), but the ordinal position in the story averages out this modulation across the entire story. Short stimuli are dominated by boundary effects but longer naturalistic stimuli are not.) This is also visible in Figure 2, in which the regression lines have roughly the same slope regardless of index.

**Table 1:** Summary of model fit for (corpus) frequency class in the time window 300-500ms from stimulus onset using all content words.

| Linear mixed model fit by maximum likelihood |  |  |  |  |
| --- | --- | --- | --- | --- |
| AIC | BIC | logLik | deviance |  |
| 2021954 | 2021996 | -1010973 | 2021946 |  |
| Scaled residuals: |  |  |  |  |
| Min | 1Q | Median | 3Q | Max |
| -33.06 | -0.53 | 0 | 0.53 | 38.43 |
| Random effects: |  |  |  |  |
| Groups | Name | Variance | Std.Dev |  |
| subj | (Intercept) | 0.10 | 0.31 |  |
| Residual |  | 130.01 | 11.40 |  |
| Number of obs: 262392, groups: subj, 52. |  |  |  |  |
| Fixed effects: |  |  |  |  |
|  | Estimate | Std. Error | t value |  |
| (Intercept) | 0.5 | 0.075 | 6.6 |  |
| corpus.freq | -0.059 | 0.0045 | -13 |  |

#### 3.1.2 *Relative Frequency*

The relative frequency of a word in a story is also known to correlate with N400 amplitude (cf. Van Petten et al., 1991, who found a repetition priming effect for words repeated in natural reading). This is seen indirectly in repetition priming (which is essentially a minimal, binary context) and information-theoretic surprisal, which can be seen as a refinement of relative frequency. In con-trast to corpus frequency, incorporating index does improve the relative frequency model (see Table S4 in the supplementary materials). The improved model is presented in Table 2; relative frequency was divided into logarithmic classes using the same algorithm as for corpus frequency, but applied exclusively to the smaller “corpus” of the story. Interestingly, the interaction of index with relative frequency has a smaller estimated value than the main effect for index, but a larger *t*-value, indicating a more reliable estimate and a clearer effect. This interaction is visible in the clearly differing slopes in Figure 3. The main effect for relative frequency has both a larger estimate and *t*-value than the terms with index.

**Figure 1:**
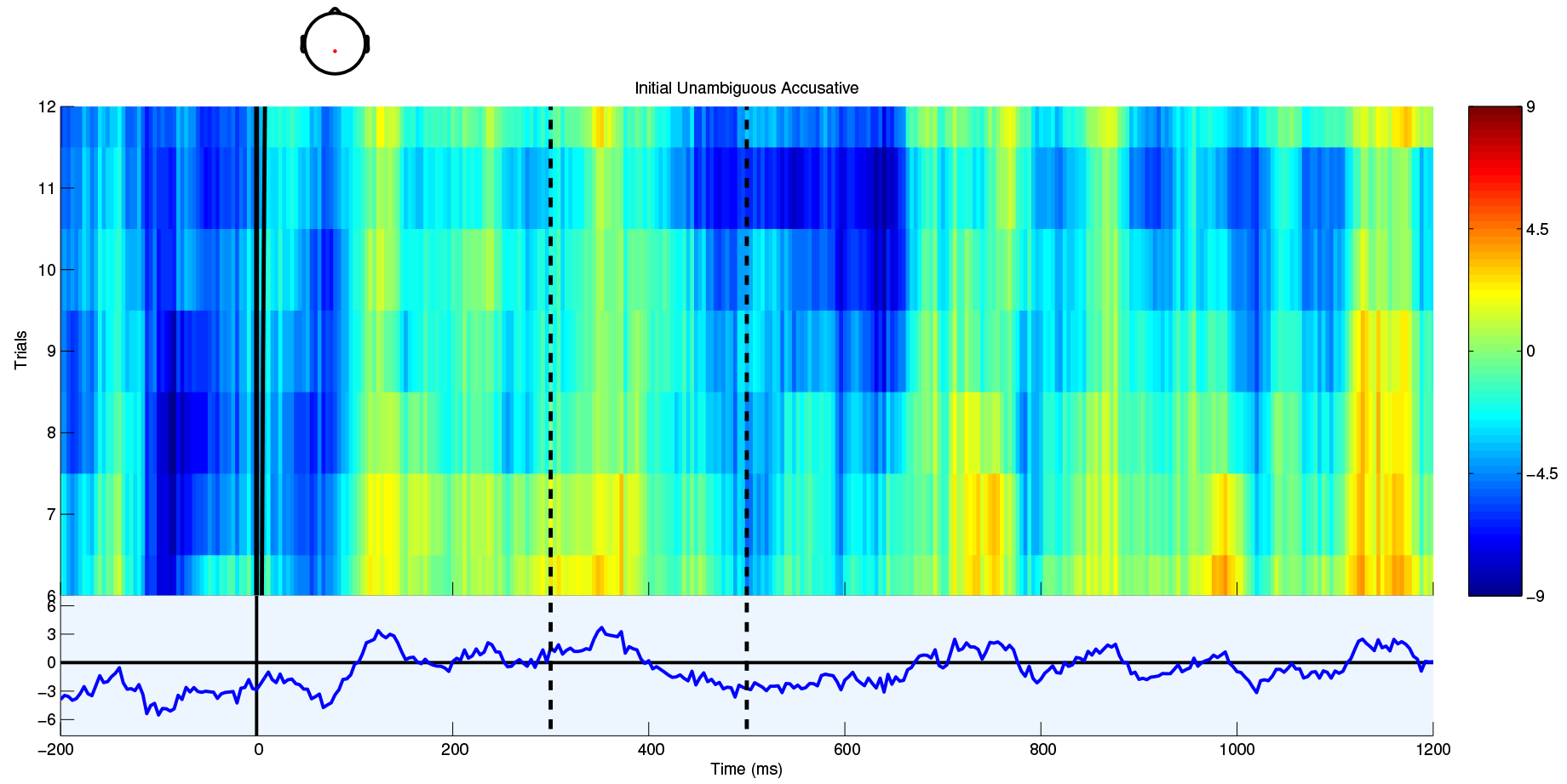
Single trial and average ERPs from electrode CPz from a single subject for unambiguous accusatives placed before a nominative. In the upper part, single trials are displayed stacked and sorted from top to bottom in decreasing orthographic length as a weak proxy for acoustic length, while the lower part displays the average ERP. Amplitude is given by color in the upper part and by the *y*-axis in the lower part. The dashed vertical lines indicate the boundaries of the N400 time window, 300 and 500ms post stimulus onset.

### 3.2 *Frequency is Dynamic*

Somewhat surprisingly, the model for relative frequency with index provides nearly as good a fit as the model for corpus frequency (Table 3). Adopting a Bayesian perspective on the role of prior information (here: frequency), this result is less puzzling. From a Bayesian perspective, corpus frequency is a nearly universally applicable but weakly informative prior on the word, while the relative frequency is (part of) a local prior on the word. This is clearly seen in the interaction with position in the story - corpus frequency’s informativeness does not improve over the course of the story, but relative frequency’s does as the probability model it represents is asymptotically approached. (This is in line with previous sentence-level findings that frequency effects are strongest early on, cf. Payne et al. (2015).) Thus, (corpus) frequency makes a small but measurable contribution in a rich context, while it tends to dominate in more restricted contexts. Relative frequency becomes a more accurate model of the world, i.e. a more informative prior, as the length of the context increases. Corpus frequency is thus in some sense an approximation of the relative frequency calculated over the context of an average speaker’s lifetime of language input.

**Figure 2:**
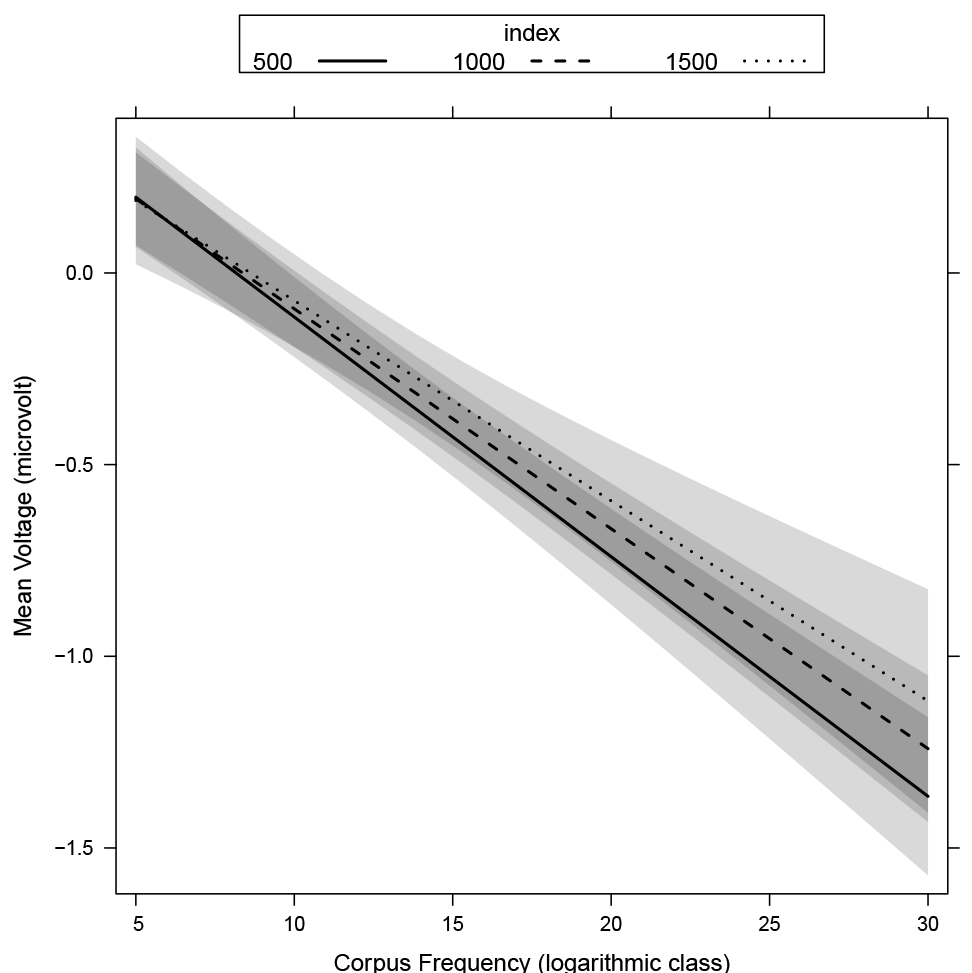
Plot of effects for corpus frequency interacting with index (ordinal position in the story). Shaded areas indicate 95% confidence intervals. Index is divided into tertiles and plotted in an overlap to make the lack of interaction more prominent. There is an increasing negativity with decreasing frequency (higher logarithmic class), which is unaffected by position in the story.

In this sense, we can say that frequency is dynamic and not a static, inherent property of a word. In the absence of local context, frequency is calculated according to the most general context available - the sum total of language input. With increasing local context, a narrower context for calculating frequency is determined, increasingly cut down from the global language input (which now of course includes the new local context). From this perspective, it is less surprising that a model incorporating the development of relative frequency over time yields results that are nearly as good as a model based on the well-established effect of corpus frequency. Frequency is an approximation for expectation, and a larger context leads to expectation that is better predicted from that context than from general
trends.

**Table 2:** Summary of model fit for relative frequency class and index (ordinal position) in the time window 300—500ms from stimulus onset using all content words. The interaction term yields a reliable estimate, while the main effect for index is not quite reliable.

| AIC | BIC | logLik | deviance |
| --- | --- | --- | --- |
| 2022079 | 2022141 | -1011033 | 2022067 |

Scaled residuals:
| Min | 1Q | Median | 3Q | Max |
| --- | --- | --- | --- | --- |
| -33.07 | -0.53 | 0 | 0.53 | 38.4 |

Random effects:
| Groups | Name | Variance | Std.Dev |
| --- | --- | --- | --- |
| subj | (Intercept) | 0.10 | 0.31 |
| Residual |  | 130.07 | 11.40 |

Number of obs: 262392, groups: subj, 52. Fixed effects:
|  | Estimate | Std. Error | t value |
| --- | --- | --- | --- |
| (Intercept) | 0.51 | 0.17 | 3 |
| index | −0.00031 | 0.00017 | −1.8 |
| rel.freq | −0.14 | 0.027 | −5.3 |
| index:rel.freq | 6.7e−05 | 2.8e−05 | 2.4 |

### 3.3 *Animacy, Case Marking and Word Order*

In addition to frequency as a relatively basic, word-level property, we examined the effects of several higher-level cues to sentence interpretation - animacy, case marking and word order - in order to determine whether our methodology is also suited to examining neural activity related to the interpretation of linguistically expressed events. Psycholinguistic studies using behavioral methods have demonstrated that such cues play an important role in determining real-time sentence interpretation (e.g. with respect to the role of a participant in the event being described; a human is a more likely event instigator, as is an entity that is mentioned early rather than late in a sentence etc.) - and, hence, expectations about upcoming parts of the stimulus (e.g. Bates et al., 1982; MacWhinney et al., 1984). Electrophysiological evidence has added support to this claim, with an increased N400 amplitude for dispreferred yet grammatically correct constructions (e.g. for accusative-initial sentences in several languages including German, Swedish and Japanese, see Schlesewsky et al. (2003); Wolff et al. (2008); Bornkessel et al. (2003); Hörberg et al. (2013); for animacy effects in English, Chinese and Tamil, see Weckerly and Kutas (1999); Bourguignon et al. (2012); Philipp et al. (2008); Muralikrishnan et al. (2015)). As a further exploration, we examine the feasibility of measuring these effects in the natural story context.

**Figure 3:**
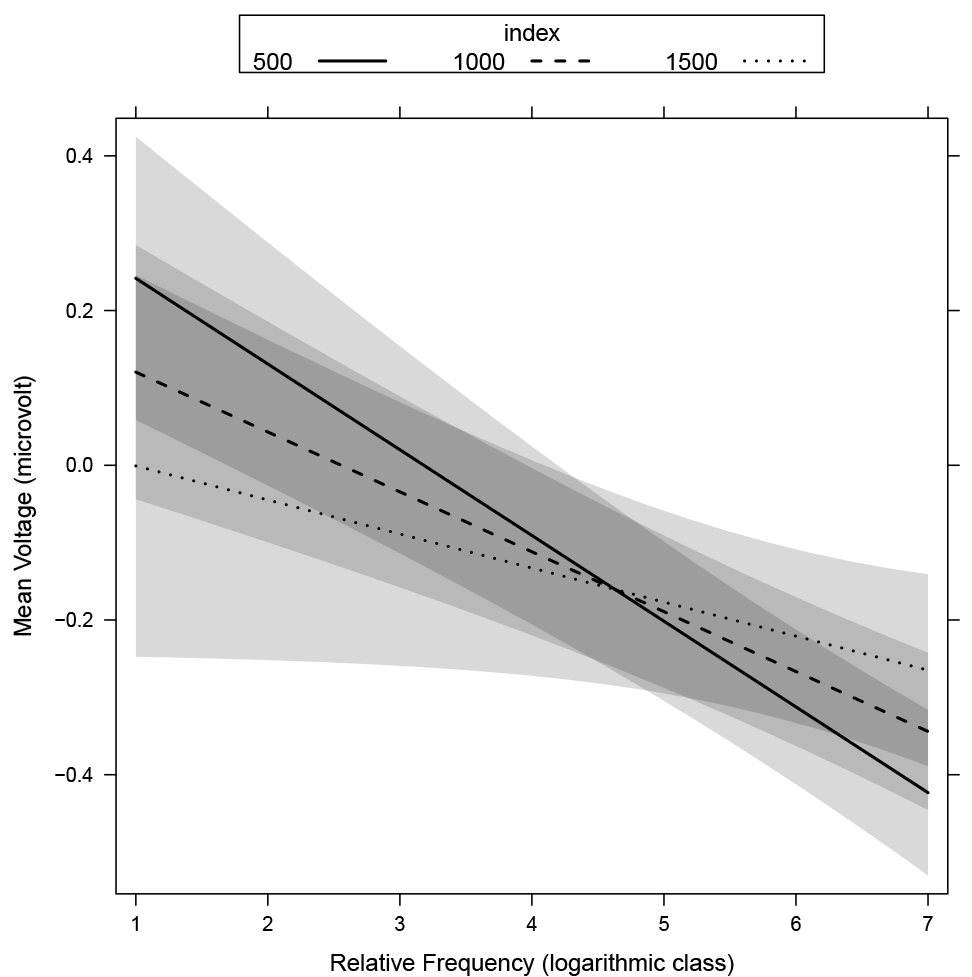
Plot of effects for relative frequency interacting with index. Shaded areas indicate 95% confidence intervals. Index is divided into tertiles and plotted in an overlap to make the interaction more prominent.

**Table 3:** Comparison of best models for corpus and relative frequency. Both models yield similar fits as evidenced by log-likelihood. The additional parameters of the relative-frequency model lead to somewhat poorer AIC and BIC values.

|  | Df | AIC | BIC | logLik |
| --- | --- | --- | --- | --- |
| m.freq | 4 | 2021954 | 2021995 | -1010973 |
| m.rel.index | 6 | 2022078 | 2022141 | -1011033 |

For the following analyses, we further restricted the trials to full noun phrases occurring as main arguments of verbs that were in the nominative or accusative case (roughly “subjects” and “objects”, not including indirect objects). This matches previous work most closely and avoids more difficult cases where the theory is not quite as developed (i.e., what is the role of animacy in prepositional phrases?). In the following, we present the contrast for dispreferred (i.e., inanimate, non-initial position, unambiguous accusative) configurations compared to the (grand) mean (i.e. sum encoding testing *main* and not *simple effects*), and the particular arrangement *dispreferred > (grand) mean* structures the model such that the contrasts align with increased N400 activity. For morphology, there is an additional neutral classification for ambiguous case marking, and there are thus two contrasts for the unambiguous cases: *accusative (dispreferred) > (grand) mean* and *nominative (preferred) > (grand) mean*.

We begin with a model for these linguistic cues and their interactions with each other, summarized with Wald tests in shown in Table 4 and shown in full in Table 5. From the model summary, we see main effects for both types both types of unambiguous case marking, with a negativity for unambiguous nominative / preferred and a positivity for unambiguous accusative / dispreferred, which at first seems to contradict previous evidence that dispreferred cue forms elicit a negativity. This somewhat surprising result is quickly explained by the interaction between morphology and position, which shows a negativity for the dispreferred initial-accusative word order. The missing main effect for animacy at first seems contrary to previous findings, but not surprising given the limited data and its involvement in higher-level interactions, and may result from imbalance in the emergent “design” in a naturalistic stimulus.

The Wald tests show similar results with the curious exception that animacy is significant. This is likely a result of the interaction terms, even though none of them achieve the |*t*| > 2 threshold individually: animacy is important for the model; however, its effect is distributed throughout its interactions. As the Wald tests are *marginal tests*, they test the effect of completely removing a given term - and thus all of its interactions - from the model. With this in mind, it becomes clear that animacy achieves significance via its interactions. Since it is problematic to interpret main effects in the presence of interactions anyway, this is not a large problem (cf. Venables, 1998).

**Table 4:** Type-II Wald tests for the model presented in Table 5

| | $F$ | Df | Df.res | $\Pr(>F)$ |
| --- | --- | --- | --- | --- |
| animacy | 6.46 | 1 | 69045 | 0.011 |
| morphology | 16.84 | 2 | 69045 | < 0.001 |
| position | 9.10 | 1 | 69045 | 0.00256 |
| animacy:morphology | 0.33 | 2 | 69045 | 0.721 |
| animacy:position | 0.11 | 1 | 69045 | 0.74 |
| morphology:position | 23.61 | 2 | 69045 | < 0.001 |
| animacy:morphology:position | 1.85 | 2 | 69045 | 0.157 |

**Table 5:** Summary of model fit for linguistic cues (animacy, morphology, linear position) known to elicit N400-like effects. Dependent variable are single-trial means in the time window 300-500ms from stimulus onset using only subjects and (direct) objects. For animacy and position, the coefficients are named for the dispreferred condition and represent the contrast “dispreferred > preferred”. Morphology also has an additional ‘neutral’ level for ambiguous case marking, and so the coefficients represent the contrast to that level. Scaled deviation (sum) encoding was used so that the coefficients are directly interpretable as the difference between means in the given contrast.

| Linear mixed model fit by maximum likelihood |  |  |  |  |
| --- | --- | --- | --- | --- |
|  | AIC | BIC | logLik | dev |
|  | 530425 | 530553 | -265199 | 5 |
| Scaled residuals: |  |  |  |  |
|  | Min | 1Q | Median |  |
|  | -11.87 | -0.54 | 0 |  |
| Random effects: |  |  |  |  |
|  | Groups | Name | Variance | Std |
|  | subj | (Intercept) | 0.20 |  |
|  | Residual |  | 125.99 |  |
| Number of obs: 69108, groups: subj, 52. |  |  |  |  |
| Fixed effects: |  |  |  |  |
|  |  | Estimate | Std. Error | t |
|  | (Intercept) | −0.38 | 0.094 |  |
|  | inanimate | −0.051 | 0.14 |  |
|  | accusative | 0.69 | 0.22 |  |
|  | nominative | −0.72 | 0.22 |  |
|  | non-initial | −0.58 | 0.14 |  |
|  | inanimate:accusative | 0.24 | 0.45 |  |
|  | inanimate:nominative | 0.23 | 0.43 |  |
|  | inanimate:non-initial | −0.028 | 0.29 | − |
|  | accusative:non-initial | 2 | 0.45 |  |
|  | nominative:non-initial | −2.9 | 0.43 |  |
|  | inanimate:accusative:non-initial | −1.5 | 0.9 |  |
|  | inanimate:nominative:non-initial | 1.6 | 0.86 |  |

### 3.4 *Index and Corpus Frequency: Covariates, not Confounds*

We also considered more extensive models with the co-variates index and corpus frequency. Including index and corpus frequency improves the model fit (see Table S8 in the supplementary materials for full comparison). Figures 4, 5 and 6 show selected effects from this model; a full summary and selected Wald tests can be found in Tables S6 and S7 in the supplementary materials.

**Figure 4:**
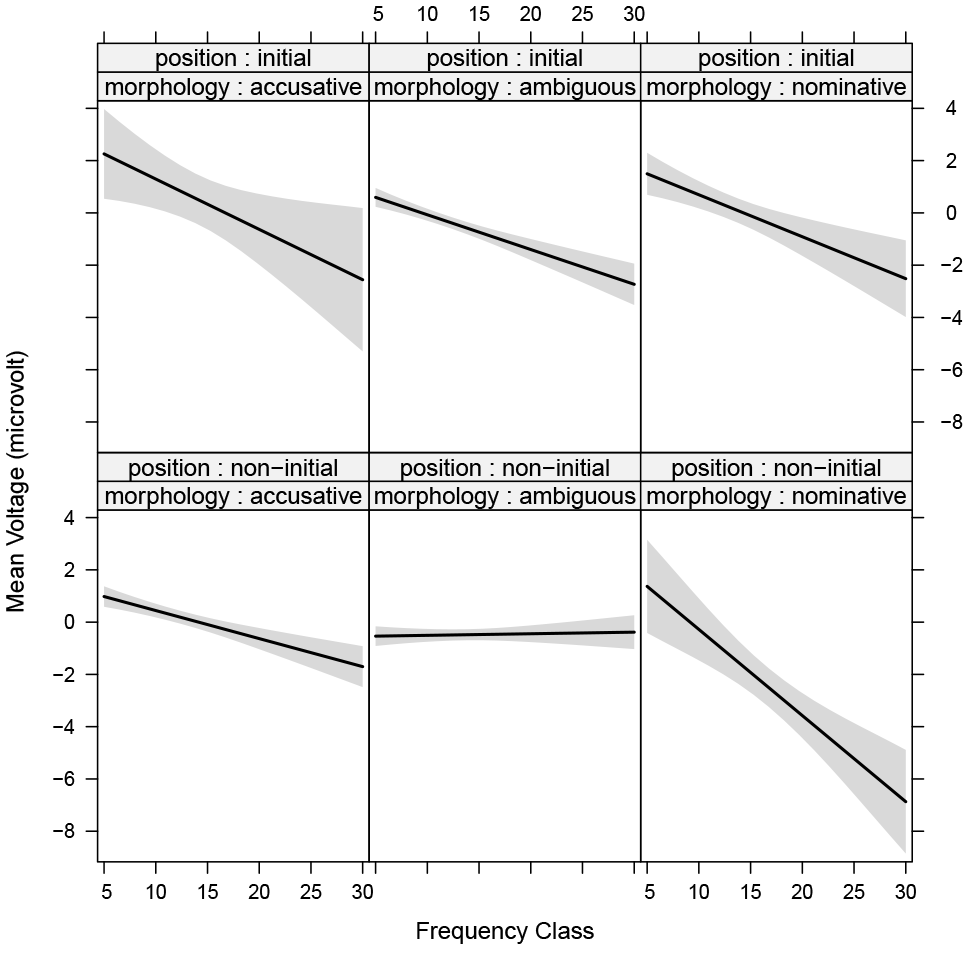
Interaction of position, morphology and corpus frequency from the full prominence model with index and frequency class. Shaded areas indicate 95% confidence intervals. Interactions with position show themselves as differences between the top and bottom rows, while interactions with morphology show themelves as differences between columns.

In the full model, we find main effects for index, corpus frequency, morphology and position. There is no main effect for animacy. This can be explained by its interactions and the reliable correlation between animacy and frequency (in this story, Kendall’s *τ* = −0.24, *p* =< 0.001), and so the variance explained by animacy is absorbed into the frequency term. The interaction between morphology and position is again present. Both morphology and position interact with frequency individually and in a three-way interaction (Figure 4). There is also a three-way interaction between the linguistic cues (Figure 5). Additionally, there are a number of higher level interactions between morphology or position, but we avoid interpreting these further than to note that they are compatible with results in the literature.

The lack of (non-nested) interaction between corpus frequency and index is also present in this model, which is visible in the level curves lying parallel to the x-axis in certain panels in Figure 6, i.e. the effect of corpus frequency does not change as a function of index. The change in patterns across panels is indicative of a higher-level interaction, i.e. there is an interaction nested within some combinations of factors, which the Wald tests bear out.

**Figure 5:**
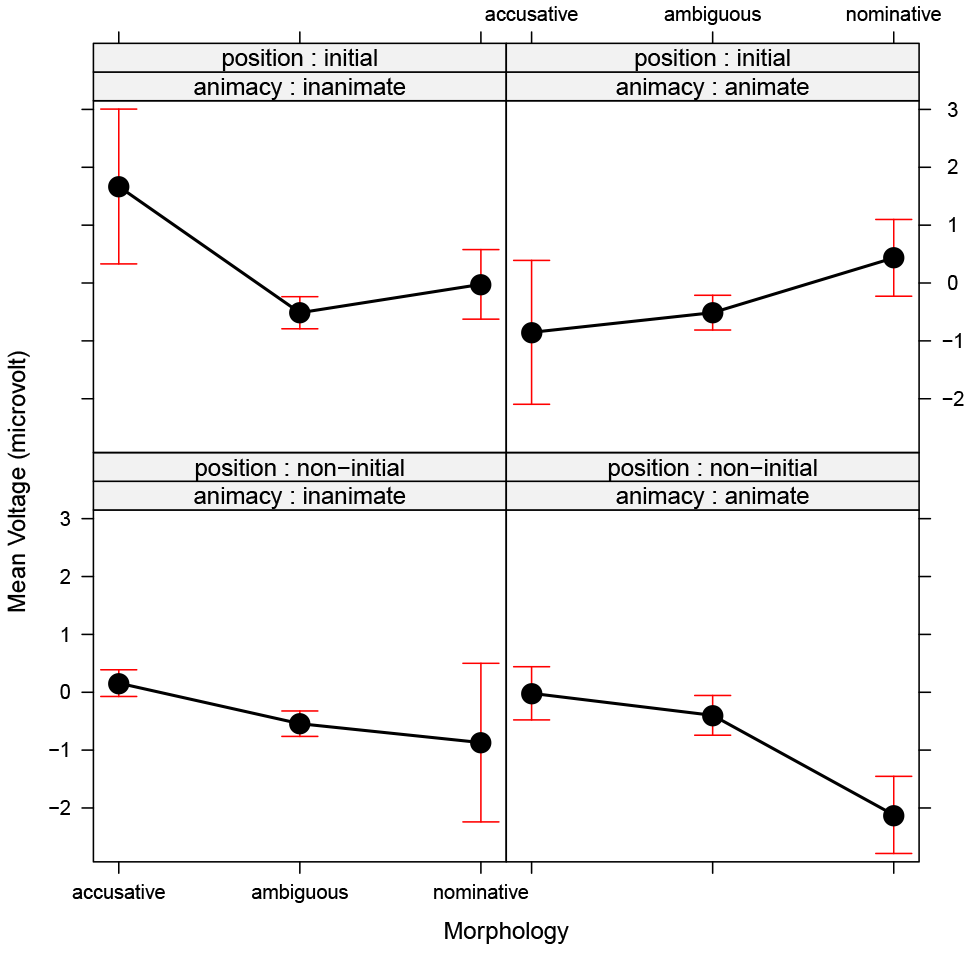
Interaction of animacy, morphology and position from the full prominence model with index and frequency class. Bars indicate 95% confidence intervals. Interactions with position show themselves as differences between the top and bottom rows, while interactions with animacy show themelves as differences between columns.

### 3.5 *Word Length*

Due to convergence issues, it was not possible to create a maximum model including orthographic length, index, corpus frequency, and all the linguistic cues, but the model with corpus frequency and orthographic length as covariates for the prominence features shows a similar set of effects (see Tables S9 and S10 in the supplementary materials). This again serves as a validity check that the effects for the linguistic cues are not merely the result of confounds with other properties of the stimulus.

**Figure 6:**
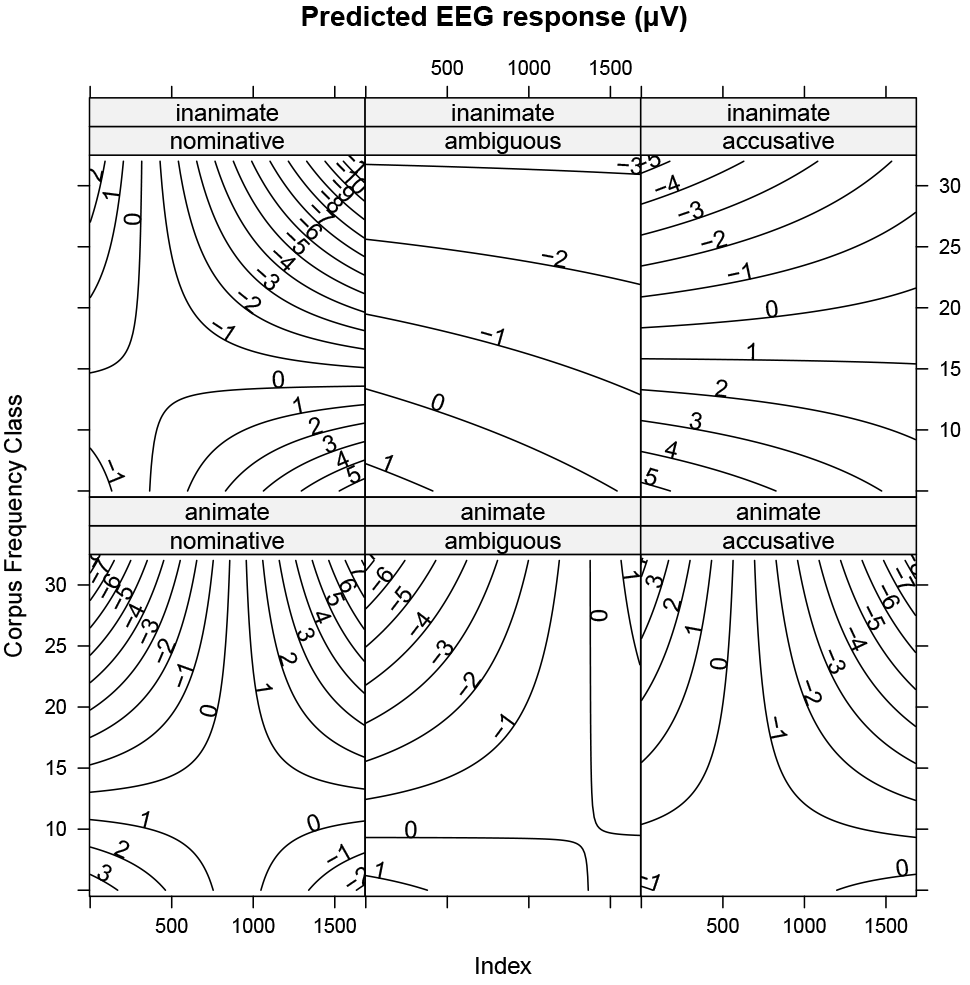
Level curves in the EEG as predicted by the full prominence model with index and frequency class. Main effects for index and frequency are indicated by change in level along the x and and y axes, respectively. Interactions between index and frequency (nested here within higher-level interactions) show themselves as “bends” in the level curves, i.e. changes in the slope in one direction along the other direction. Interactions between prominence features (animacy, morphology) and quantitative measures show themselves as different patterns of level curves across the subpanels. In particular, differences between the top and bottom row indicate an interaction with animacy and differences between columns indicate an interaction with morphology. The difference across all panels is indicative of a four-way interaction, which can also be found in the Wald tests (see Table S6 in the supplementary materials).

**Figure 7:**
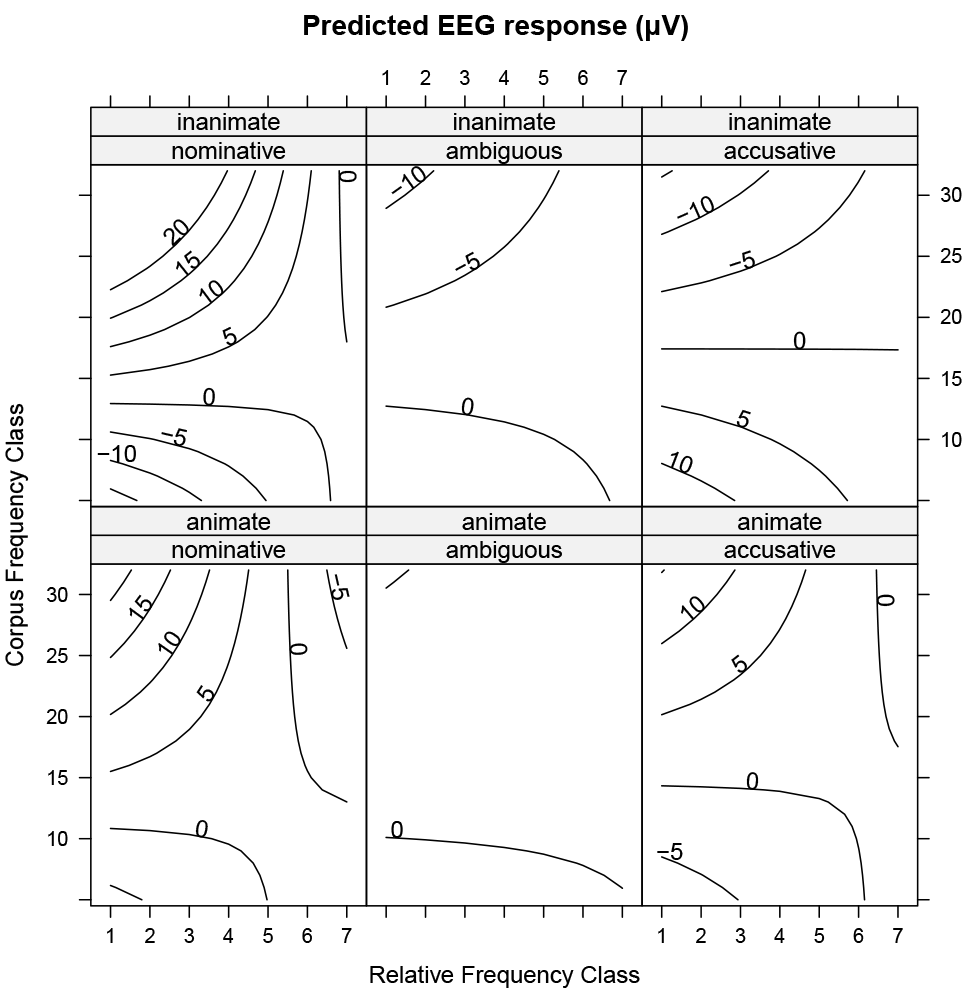
Level curves in the EEG as predicted by the combined frequency model with prominence. Main effects for relative and corpus frequency are indicated by change in level along the *x* and and *y* axes, respectively. Interactions between relative and corpus frequency show themselves as “bends” in the level curves, i.e. changes in the slope in one direction along the other direction. Interactions with animacy show themselves as differences between the top and bottom rows, while interactions with morphology show themselves as differences between columns. In highly informative or marked contexts, e.g. inanimate nominatives, the effect of frequency is completed dominated by local information and the negativity associated with decreasing frequency (i.e. increasing logarithmic frequency class) disappears or is even reversed. This is seen in the flattening or even reversal of slope of the level curves across panels.

### 3.6 *Frequency is Dynamic, Redux*

We can also examine the interplay between linguistic cues and the two types of frequency in a single model, plotted with level curves in Figure 7 (see Tables S11 and S12 supplementary materials for full model summary and selected Wald tests). Due to convergence issues, it was not possible to include index or orthographic length in this model, but nonetheless several interesting patterns emerge.

There are main effects for both types of frequency as well as morphology; additionally corpus and relative frequency interact with each other. The interaction between morphology and position is again present as well as an interaction between animacy and morphology and a threeway interaction between all three features. Interestingly, there appears to be a division in the interactions between linguistic cues and frequency type. Corpus frequency interacts with position, morphology, and with both in a three-way interaction, while relative frequency interacts with animacy and with animacy and morphology and with morphology and position in three-way interactions. There are also higher-order interactions including both frequency types and the prominence features.

## 4 General Discussion

### 4.1 *The present approach: examining complex influences within a fixed epoch*

The results for frequency in both its forms are not surprising in the sense that they match previous results. Nonetheless, it is perhaps somewhat surprising that it is possible to extract the effects in such a heterogeneous and noisy environment. Part of the problem with the type of presentation in Figure 1 is that the influences on N400 ( and, more generally, ERP) amplitude are many, including frequency, and this three dimensional representation (time on the *x*-axis, trial number sorted by orthographic length on the *y*-axis, and amplitude as color, or equivalently, on the *z*-axis) shows only some of them. Some hint of this complexity is visible in the trends between trials - the limited coherence of vertical stripes across trials reflects the sorting according to orthographic length. Unsorted, the stripes are greatly diminished. Similarly, other patterns emerge when we (simultaneously) sort by other variables, but our ability to represent more dimensions graphically is restricted.

A further complication is the inclusion of continuous predictors. Traditional graphical displays - and statistical techniques - are best suited for categorical predictors, which we can encode with different colors, line types or even subplots. However, the mixed-effects models are capable of incorporating many dimensions simultaneously, including continuous dimensions like frequency, which have been traditionally difficult to present as an ERP without resorting to methods like dichotomization (see Smith and Kutas, 2014a; Smith and Kutas, 2014b, for a similar but complementary approach using continuous-time regression; see Payne et al., 2015, for a similar approach at the sentence level for a continuous-measure reanalysis of an older, dichotomously analyzed study). In other words, traditional graphical representations of ERPs have difficulty displaying more complex effects and interactions.

Our approach is to pick a fixed time-window, freeing up the horizontal axis for something other than time, which fits well with the epoch-based regression approach used here and in Payne et al. (2015). Displays of the regression at a particular time point are also level curves at a particular time and provide clarity about the shape of the effect at a particular time, but are less useful for exploring the time course of the ERP. Nonetheless, this perspective allows us to study the modulation of the ERP in a given epoch via more complex influences, such as those that arise in a natural story context. The implications of this perspective - complex influences in a fixed epoch - are discussed more645 fully below.

### 4.2 *Implications for Electrophysiological Research in Cognitive Neuroscience: ERP Components as Ongoing Processes*

Thus far, we demonstrated that the synthesis of increasingly tractable computational techniques (mixed-effects models, automatic artefact correction with independent-component analysis) leads to a tractable approach to analyzing electrophysiological data collected in response to a naturalistic auditory stimulus (a natural story). Strikingly, the current results mirror a number of well-established findings from traditional, highly controlled studies. This is somewhat surprising given the large amount of jitter in naturalistic stimuli. The words themselves have different lengths and different phonological and acoustic features; moreover, the phrases have different lengths, which are often longer than in traditional experiments. This leads to the information carried by the acoustic-phonological signal being more broadly and unevenly distributed in time. Yet, we still see clear effects at a fixed latency, which seems to be at odds with traditional notions of ERPs as successive, if occasionally overlapping events (i.e. components), reflecting various (perhaps somewhat parallel) processing stages. In the following, we discuss the implications of our results for the interpretation of ERP responses in cognitive neuroscience research - both in a naturalistic auditory environment and beyond.

From the traditional perspective - that ERPs are the sum of discrete components - individual components within the electrophysiological signal (e.g. the N200, N400, P300 and P600 to name just a small selection of examples) are interpreted as indexing particular cognitive processes which occur at certain, clearly defined times within the overall time course of processing (see e.g. Friederici, 2011, for a recent review in the language domain). However, ERP data recorded in response to naturalistic, auditory language challenge this traditional view: in contrast to ERPs in studies employing segmented visual presentation (RSVP), components no longer appear as well-defined peaks during ongoing auditory stimulation and this applies equally to the early exogenous components and to endogenous components.

Let us first consider the exogenous components. The fact that these no longer appear during continuous auditory stimulation other than at stimulus onset does not mean that the neurocognitive processes indexed by these early components do not take place later in the stimulus, but rather that their form is no longer abrupt enough to be visually distinct from other signals in the EEG. The abruptness of stimulus presentation in RSVP leads to the abruptness of the components, but continuous stimulation, as in a naturalistic paradigm, leads to a continuous modulation of the ERP waveform without the typical peaks of RSVP.

More precisely, the relevant continuity is not that of the stimulus itself, but rather of the information it carries. In RSVP, *all* external information for a given presentation unit is immediately available, although there may be certain latencies involved in processing this information and connecting to other sources of information (e.g. binding together multimodal aspects of conceptual knowledge). Thus, as the information passes through the processing system, it is available in its entirety and there are sharp increases in neural activity corresponding to this flood of new information resulting in sharp peaks. In auditory presentation, the amount of external information is transmitted over time (instead of over space), and thus the clear peaks fall away as the incoming information percolates continuously through the processing system, yielding smaller and temporally less well-defined modulations of the ERP. In summary, we propose that the appearance of ERP components as small modulations or large peaks is a result of the relative change in the degree of information processed. In studies employing visual presentation, time-locking to recognition point (e.g. van der Brink and Hagoort, 2004; Wolff et al., 2008) or employing other similar jitter-controlling measures in auditory presentation, ERPs thus reflect the state of processing *at the climax of (local) information input* and fail to provide information about incrementality below the level of units such as words.

This proposal for extreme incrementality accords well with a predictive coding-based approach to electrophysiological responses, in which ERP responses such as the mismatch negativity (MMN) reflect both bottom-up adaptation to the stimulus and modulation of top-down predictions / adjustment of an internal model (Friston, 2005; Garrido et al., 2009). Predictive coding posits that the brain constantly attempts to match sensory input sampled from the external world to predictions about the state of the world derived from an internal model, accomplished by means of hierarchically organized forward and inverse models and thought to be implemented by hierarchically organised cortical networks. At the lowest level, predictions are matched against sensory input and any resulting mismatch (prediction error) is propagated back up the hierarchy via feedforward connections (bottom-up adaptation), thereby initiating model updates to minimise prediction errors both at the current level and the level below (top-down modulation). From the predictive coding perspective, the MMN for deviant stimuli within a series of standards reflects an attenuation of the response to the standards rather than the generation of an additional mismatch response to the deviants: stimulus repetition leads to model adjustment and the minimization of prediction error for subsequent standard presentations and, accordingly, a disappearance of the MMN. An approach along these lines straightforwardly accounts for the apparent discrepancy between ERP responses in traditional and naturalistic paradigms. In naturalistic settings, continuous stimulation in conjunction with rich contextual information leads to increased model update and adaptation, particularly for early sensory aspects of processing, thereby resulting in an attenuation of ERP components. In other words, the prediction errors and resulting model updates are necessarily more pronounced in isolated stimuli than in stimuli encountered in a naturalistic context. While our approach does not directly model the low-level neural computations of the predictive-coding framework (for that, see e.g. Friston et al., 2012a), it does show that its explanatory framework provides for a coherent, parsimonious account of EEG/ERP activity during naturalistic stimulation.

### 4.3 *Continuous Components, Continuous Processing, and Growing Representations*

We propose that this continuous, subsymbolic incrementality can be extended to also account for a broader range of stimulus-locked components such as the N200 and N400. Specifically, we suggest that the account of the MMN outlined above can be straightforwardly extended to these components in the sense that they reflect similar stimulus-related processing mechanisms as the MMN (bottom-up adaptation and top-down modulation), but at different levels of the processing hierarchy (for a somewhat similar view, see Pulvermüller et al., 2009). This view is not entirely new: early research concerning the N400 examined the possibility that it was a member of the N200 family (Kutas and Federmeier, 2011), much like the long-standing debate about whether the P600 belongs to the P300 family (e.g. Gunter et al., 1997; Osterhout et al., 1996; Coulson et al., 1998; Sassenhagen et al., 2014). The notion of continuous processing presented here hints at a coherent account for such component families, related to their temporal resolution. Following Giraud and Poeppel (2012)’s suggestion that the frequency bands in cortical oscillations track the time resolution of hierarchical structure in speech processing, we can consider similar ERP components with different time-scales as tracking the time resolution of different stimulus features (see also Bornkessel and Schlesewsky, 2006; Dogil et al., 2004; Roehm et al., 2007; cf. “temporal receptive windows”, Hasson et al., 2008; Lerne et al., 2011; see also Henry and Obleser, 2012; Herrmann et al., 2016, for frequency-band entrainment). In this view, the MMN and N200 are similar to the N400 but react to more basic features of the stimulus at a lower latency because they reflect a similar neural process earlier in the processing hierarchy. This leads to a higher temporal resolution but a smaller analysis time window, in accordance with the frequency of the oscillation under consideration. This perspective accounts for the apparent paradox of MMN effects for manipulations more typical to N400 experiments (cf. “ultrafast processing” in recent studies such as Pulvermüller et al., 2001; MacGregor et al., 2012; Shtyrov et al., 2014); or other fast recognitions of large scale stimulus change (e.g. category error in Dikker et al., 2009) as reflecting predictions that are exceedingly precise and can thus be falsified quickly. Moreover, similar mechanisms operating at different scales is compatible with the recent proposal that the mechanisms for human language processing arise from a difference from nonhuman primates in quantity rather than quality (Bornkessel-Schlesewsky et al., 2015) and is compatible with the account that the neural aspects of early language acquisition follow increasing time scales (Friederici, 2005). More complex processing arises as fundamental processing mechanisms are repeated and expanded across multiple time scales.

Taken together with O’Connell et al. (2012)’s “continuous oddball” design, our results suggest an answer to some of the outsanding question posed by Hasson et al. (2015), namely the continuity of information transmission in process memory and the relationship between process memory and information integration during decision making. Information is passed continuously along the processing hierarchy, but bursts may occur based on discontinuities in the stimulus or emergent properties of processing (exceeding a given threshold; accumulation of evidence leading to a tipping point, cf. Rousselet and Pernet, 2011, who suggest that peaks may reflect outputs and not mechanisms themselves). In the case of stimulus-locked components, the time-course of processing is matched to properties of the stimulus, even responding to the the compression and dilation of the input (Lerner et al., 2014). However, even this dynamic adaption has limits, which may be linked to intrinsic properties of neural computation, - exceeding these limits leads to a breakdown in both the temporal scaling of processing and intelligibility (Lerner et al., 2014). In the case of response-locked components, peak-like behavior reflects thresholded behavior, e.g. binary decision making, but slow drifts during the accumulation phase are possible. The smearing we observed here for an epoch-based approach for the N400 has also been visible for years in traditional stimulus-locked analyses of the P600, a response-locked component (Sassenhagen et al., 2014). The broad, slow positive waved observed in traditional stimulus-locked ERP analyses results from the smearing induced by variable response-latencies, which is equivalent to sampling the component at various stages in the response process, i.e. the response complement to the perception problem considered here. In brief, information processing is continuous, but may exhibit emergent “pulsatile” behavior based on thresholding and uneven distribution of information in the stimulus. As such words and other “atoms” of communication are emergent phenomena based on stable configurations in the temporal distribution of the communication signal, whether phonological, orthographic or gestural (cf. “attractor basins” in much of the literature; for a predictive-coding perspective, see Friston et al. (2012b) and Alday et al. (2014) for an application to language).

## 5. Conclusion

We have demonstrated the feasibility of studying the electrophysiology of speech processing with a naturalistic stimulus through a synthesis of modern computational techniques. The replication of well-known effects serve as a proof of concept, while initial exploration of the more complex interactions possible in a rich context suggested new courses of study. Surprisingly, we found robust manipulations at a fixed latency from stimulus onset in spite of the extreme jitter from differences in word and phrase length. This suggests that ERP responses should be viewed as continuous modulations and not discrete, yet overlapping waveforms.

## Acknowledgements

We would like to thank Fritzi Milde for her help in annotating the stimulus, Jon Brennan for helpful discussions related to naturalistic stimulus presentation and EEG/MEG measures, Jona Sassenhagen and Franziska Kretzschmar for engaging discussions, and Jona Sassenhagen again for his help with EEGLAB.

## Footnotes

1 While it is possible to calculate effect sizes, etc. from ANOVA results, this is generally a post-hoc test and not delivered by the ANOVA procedure directly. Moreover, mixed-effects models estimate *parameters* in a quantitative model framework directly, and not just *effect sizes*, and accomodate shrinkage and other issues related to the Stein paradox (Efron and Morris, 1977; Stein, 1956), which simple summary statistics like the grand mean do not do.

2 For larger number of electrodes, it is possible to include a random-effect term for “electrode” (cf. Payne et al., 2015); however, this can be problematic. There is systematic, parametric variation between channels (topography) that is perhaps best modelled as a fixed-effect (e.g as “sagitality” or “laterality”). If channels without respect to topography are modelled as a random effect, then we can fail to capture this parametric variation; moreover, this variation may violate assumptions about the (multivariate normal) distribution of the random effects. If channels are modelled as a random effect constrained by topography (e.g. within ROIs), then low-density electrode configurations, such as the one used here, will not provide enough levels to accurately model the random effect: random effects are variance components (see supplementary materials), and are thus, like all estimates of variance, extremely sensitive to small sample sizes due to their skewed distribution.

3 For a brief overview, see Dale Barr’s explanation online at http://talklab.psy.gla.ac.uk/tvw/catpred/.

4 The converse arrangment *preferred* > (*grand*) *mean* would yield a model with coefficients indexing *decreased* N400 activity.

5 Indeed, this is how (marginal) — tests differ from Type-I (sequential) and Type-III (non-sequential, i.e. not sensitive to order and keeping higher-order interactions when testing a lower-order term).

6 While modern ERP theories do not assume discrete events and thus easily allow for continuous modulation, the common intuition seems to be based on a weak-form of *ERPology* (cf. Luck, 2005) with discrete, if overlapping, components.

